# Parkinson’s Paradox: Alpha-synuclein’s Selective Strike on SNc Dopamine Neurons over VTA

**DOI:** 10.1101/2025.03.24.644952

**Authors:** L Phan, D Miller, A Gopinath, M Lin, D Gunther, K Kiel, S Quintin, D Borg, Z Hasanpour-Segherlou, A Newman, Z Sorrentino, J Miller E, J Seibold, B Hoh, B Giasson, H Khoshbouei

## Abstract

In synucleinopathies, including Parkinson’s disease (PD), dopamine neurons in the substantia nigra pars compacta (SNc) exhibit greater vulnerability to degeneration than those in the ventral tegmental area (VTA). While α-synuclein (αSyn) pathology is implicated in nigral dopamine neuron loss, the mechanisms by which αSyn affects neuronal activity and midbrain dopamine network connectivity prior to cell death remain unclear. This study tested the hypothesis that elevated αSyn expression induces pathophysiological changes in firing activity and disrupts network connectivity dynamics of dopamine neurons before neuronal loss. We employed two mouse models of synucleinopathy: preformed αSyn fibril (PFF) injection and AAV-mediated expression of human αSyn (hαSyn) under the control of the tyrosine hydroxylase (TH) promoter, both targeting the VTA and SNc. Four weeks post-injection, brain sections underwent histological, electrophysiological, and network analyses. Immunohistochemistry for TH, hαSyn, and phospho-Ser129 αSyn assessed αSyn expression and dopaminergic neuron alterations. Neuronal viability was evaluated using two complementary approaches: quantification of TH^+^ or FOX3^+^ and TUNEL labeling. Importantly, these analyses revealed no significant changes in neuronal counts or TUNEL^+^ cells at this time point, confirming that subsequent functional assessments captured pre-neurodegenerative, αSyn-induced alterations rather than late-stage neurodegeneration. Electrophysiological recordings revealed a differential effect of hαSyn expression. SNc dopamine neurons exhibited significantly increased *baseline* firing rates, whereas VTA dopamine neurons remained unchanged. These findings indicate a region-specific vulnerability to αSyn-induced hyperactivity of dopamine neurons. Further analysis revealed impaired homeostatic firing rate regulation in SNc, but not VTA, dopamine neurons, demonstrated by a reduced capacity to recover baseline firing following hyperpolarization. Collectively, our results demonstrate that, prior to neurodegeneration, elevated αSyn expression differentially disrupts both basal firing activity and network stability of SNc dopamine neurons, while sparing VTA dopamine neurons. By identifying neurophysiological changes preceding dopaminergic neuron loss, these findings provide critical insights into the pathophysiological mechanisms predisposing SNc neurons to degeneration in Parkinson’s disease.

**Significance Statement:** A central question in Parkinson’s disease research is why dopamine neurons in the substantia nigra pars compacta (SNc) are more vulnerable than those in the ventral tegmental area (VTA). This study reveals that alpha-synuclein (αSyn) pathology differentially impacts dopamine neuronal activity and network connectivity, causing changes in the SNc before neuronal loss occurs, but not in the VTA. These findings provide a mechanism to explain the differential resilience of these neighboring dopamine neuron populations and provide insights into Parkinson’s disease progression. The methodologies developed in this study establish a foundation for investigating network topology in deep brain structures and its role in neurodegenerative disorders.

## Introduction

Parkinson’s disease (PD) is neuropathologically characterized by two defining hallmarks: (1) the presence of α-synuclein (αSyn)-containing Lewy bodies and Lewy neurites, and (2) a significant loss of dopaminergic neurons in the substantia nigra pars compacta (SNc)(Giasson and Lee 2003, Braak et al. 2003a, Stoker and Greenland 2018). Although neither hallmark is exclusive to PD, their co-occurrence is required for a definitive diagnosis. Missense mutations and duplication/triplication in the *SNCA* gene, encoding αSyn, results in PD and the related disorder dementia with Lewy bodies, strongly implicating that that αSyn pathobiology is a crucial factor in disease pathogenesis.

Alpha-synuclein is a small, natively unfolded protein that associates with curved membranes, particularly synaptic vesicles, where it modulates neurotransmitter release (Gao et al. 2023, Burré 2015, Somayaji et al. 2020). In a disease state, αSyn misfolds and aggregates into oligomeric and fibrillar forms, contributing to dopaminergic neuron dysfunction and, ultimately, cell death(Luk et al. 2012, Dryanovski et al. 2013, Chen and Feany 2005, Cookson 2009). These aberrant αSyn conformations disrupt multiple intracellular processes, including vesicle trafficking, synaptic transmission, neuronal firing, and mitochondrial function (Cookson 2009, Phan et al. 2017, Garcia-Reitböck et al. 2010, Tozzi et al. 2021, Geibl et al. 2024). Furthermore, αSyn aggregates can propagate between neurons via several proposed mechanisms, including calcium-dependent pathways, potentially spreading pathology throughout the brain (Lautenschläger et al. 2018, Emmanouilidou et al. 2010). αSyn’s role in dopamine signaling is complex, exhibiting both facilitatory and inhibitory effects that depend on neuronal activity, the specific molecular targets engaged, and the duration of αSyn deposition (Somayaji et al. 2020). Previous studies have shown that, prior to neurodegeneration, excess αSyn impairs dopamine transporter (DAT)-mediated dopamine clearance and promotes DAT-dependent, non-vesicular dopamine efflux, leading to increased extracellular dopamine levels (Swant et al. 2011, Butler et al. 2015, Lam et al. 2011). Beyond dopamine homeostasis, αSyn overexpression disrupts the intrinsic firing properties of dopamine neurons and alters their calcium dynamics (Tozzi et al. 2021, Dagra et al. 2021, Lin et al. 2021). These perturbations are hypothesized to destabilize neuronal energy balance and increase neuronal susceptibility to degeneration (Ganjam et al. 2019, Geibl et al. 2024, Ordonez, Lee and Feany 2018). Notably, αSyn pathology and associated toxicity are more pronounced in the SNc, which is highly susceptible to dopaminergic neuron loss, compared to the relatively resilient, adjacent ventral tegmental area (VTA).

A critical question in PD research is why SNc dopamine neurons are selectively vulnerable compared to VTA neurons. SNc neurons possess a distinct vulnerability profile, including αSyn-reduced TH expression, greater dendritic arborization (and thus higher metabolic demands), extensive axonal projections, and differential expression of proteins such as calbindin, L-type Ca^2+^ channels, and vesicular glutamate transporter 2 (VGLUT2) (Lebowitz and Khoshbouei 2020, Maingay et al. 2006, Pacelli et al. 2015, Braak et al. 2003b, Poulin et al. 2014, Chan, Gertler and Surmeier 2010, Steinkellner et al. 2018). Our previous work, using primary cultures of dopamine neurons overexpressing αSyn and human iPSC-derived dopamine neurons with *SNCA* triplications, demonstrated that αSyn overexpression alters dopamine neuron firing patterns and dopamine release (Dagra et al. 2021, Lin et al. 2021). However, the differential effects of αSyn deposition on the intrinsic activity of SNc *versus* VTA dopamine neurons within an intact midbrain circuit remain poorly understood. Elucidating these pre-degenerative changes in dopaminergic neuronal activity and network connectivity in these adjacent dopaminergic nuclei is crucial for developing therapeutic strategies to slow disease progression and restore neuronal function and resilience.

In this study, we found no evidence of cell loss four weeks after viral-mediated αSyn overexpression or preformed fibril (PFF) deposition in the SNc and VTA. This indicates that we captured an early, pre-degenerative stage of αSyn deposition in dopamine neurons, prior to the onset of neurodegeneration. We therefore leveraged this model to examine the effects of αSyn deposition on the intrinsic electrophysiological properties of SNc and VTA dopamine neurons, their resilience to hyperpolarization-induced challenge, and the connectivity of their dopaminergic networks. Our data demonstrate that both αSyn overexpression and PFF deposition alter the electrophysiological and network connectivity of SNc dopamine neurons, significantly impairing their capacity to maintain homeostatic function, while sparing VTA dopamine neurons. These findings shed light on the early pathogenic mechanisms contributing to SNc neuron vulnerability in synucleinopathies and provide a foundation for developing interventions aimed at preventing or slowing neurodegeneration.

## Materials and Methods

Animals: DAT^IREScre^ and Ai95(RCL-GCaMP6f)-D knock-in mice were obtained from Jackson Laboratory (Stock number: 006660 (DAT^IRES^cre), 024105 (Ai95D), Bar Harbor, ME, USA). DAT:cre, DAT:cre-GCaMP6f, and C57Bl/6 mice aged between 1.5 and 2 months and averaging 22-24 grams in weight, were utilized for this study. The mice were maintained in a controlled environment at the University of Florida’s animal care facility, with a 12-hour light-dark cycle and ad libitum food/water access. The care and experimental procedures involving these animals were conducted in adherence to the guidelines approved by the University of Florida’s Institutional Animal Care and Use Committee (IACUC).

## Recombinant αSyn proteins and preparation of αSyn preformed fibrils (PFFs)

Recombinant mouse αSyn protein was expressed in BL21 (DE3) E. coli (New England Biolabs Inc) using the pRK172 bacterial expression vector cloned with the mouse αS cDNA (Giasson et al. 2001). High salt (750 mM NaCl, 50 mM Tris, pH 7.5, 1 mM EDTA) and heat-resistant bacterial lysates were purified utilizing size exclusion chromatography followed by Mono Q anion exchange chromatography as described previously (Giasson et al. 2001, Greenbaum et al. 2005). Protein concentrations were assessed via BCA. For the generation of fibrils, mouse αSyn protein (5 mg/ml) was incubated in sterile PBS (Life Technologies) at 37 °C with continuous shaking at 1050 rpm for 5 days (Thermomixer C, Eppendorf), and fibril formation was assessed by turbidity and K114 fluorometry (Crystal et al. 2003). Fibrils were diluted to 2 mg/ml in sterile PBS and sonicated for 60 min in a water bath sonicator and stored at-80 °C until use (Waxman and Giasson 2010).

### Stereotaxic viral injection

Unless otherwise specified, 6-to 7-week-old mice were used for intracranial injections of PFF (2 µg/µL) or AAV1-TH-human-αSyn (hαSyn) virus (1 µL) in all experiments. Following anesthesia induction at 3% isoflurane, the animal was shaved, placed in the stereotaxic frame, and maintained at 1.3-1.8% isoflurane throughout surgery, with body temperature maintained at 37°C. Ophthalmic ointment was applied to prevent the eyes from drying. The surgical area was disinfected with alternating chlorhexidine followed by sterile saline. A 1 cm incision was made, and the skin was moved aside to expose lambda and bregma. The cranium was leveled to within 0.05 mm between lambda and bregma (Paxinos and Watson, 1997). At the appropriate coordinates relative to bregma a 0.5 mm burr hole was made, the dura was punctured using a scalpel and irrigated in the case of superficial bleeding. A glass micropipette backfilled with mineral oil was attached to a Drummond Nanoject III nanoliter injector and fixed to the stereotaxic apparatus. 1μL AAV1-TH-αSyn (4.92 x10^13^ U/μL) was aspirated into the glass micropipette, guided to the injection coordinates (ML = + 1.4 mm; AP = - 3.07 mm; DV = - 4.5 mm). 1µL of PFF and was injected into the SNc (ML = + 1.4 mm; AP = - 3.07 mm; DV = - 4.5 mm) and the VTA (ML = +/- 0.4 mm; AP =-3.4 mm; DV =-4.4 mm) for a total of 2 µg/µL PFF. The nanoject remained at the injection site for 10 minutes before being removed. Sterile saline was applied to the surface of the cranium and burr hole to avoid drying throughout the surgery. Postoperative analgesia (20 mg/kg Meloxicam) and hydration (1 ml sterile injection saline 0.9 %) were provided via subcutaneous injection. For 72 hours, postoperative care mice were monitored daily for body weight and provided with analgesia as recommended by veterinary staff.

### Histological validation AAV1-TH-hαSyn virus and PFF depositions

For all histological analyses, C57Bl/6 mice were microinjected with either AAV1-TH-human-αSyn (hαSyn, 1 µL) or PFF (2 µg/µL). Four weeks post-injection, mice were perfused with 1X PBS followed by 4% paraformaldehyde (PFA) in 1X PBS. Brains were dissected, stored in 4% PFA for 24–48 hours, and then transferred to 1X PBS until sectioning. Coronal midbrain sections (40 µm thick) were prepared using a Vibratome (VT 100, Leica) and stained using a free-floating protocol. Sections were first incubated in blocking solution 1 (5% fetal bovine serum (FBS) and 5% bovine serum albumin (BSA) in 0.3% Triton X-100 in 1X PBS) at 37°C for 1 hour (Table 1). They were then incubated overnight at 4°C with primary antibodies diluted in the same blocking solution (Table 2). After incubation, sections were washed four times (20 minutes each) with blocking solution 2 (5% FBS and 5% BSA in 1X PBS). They were subsequently incubated with secondary antibodies diluted in blocking solution for 1 hour at room temperature in the dark (See Table 1). Sections were counterstained with 4′,6-diamidino-2-phenylindole (DAPI). Following final washes, sections were mounted and cover-slipped using Fluoromount-G (SouthernBiotech). Images were captured using a Nikon A1 laser-scanning confocal microscope (Nikon Instruments, Melville, NY) with a Plan Fluor 20× 0.75 NA objective. Z-stack images were acquired at 1 µm intervals throughout the specimen, and maximum intensity projection images were generated. Nikon Elements NIS Analysis software was used for data acquisition.

**Table 1:** Reagents List.

| Table 1: Reagents List |  |  |  |
| --- | --- | --- | --- |
| General reagents | Concentration | Vendor | Catalog number |
| Paraformaldehyde | 4% | Electron Microscopy Sciences | 157-4-100 |
| Preformed Fibrils | 2μg/μL | Provided by Giasson Lab | N/A |
| AAV1-TH-hSyn | 4.92 x 10<sup>13</sup> U/μL | Addgene | Giasson Lab |
| AAV-FLEX-GCaMP8f | 1.1 x 10<sup>13</sup> | Addgene | 162379-AAV5 |
| **Blocking solution 1** |  |  |  |
| Triton X-100 | 0.3% | Thermo Fisher Scientific | 9002-93-1 |
| Bovine Serum Albumin | 5% | Sigma | A2153-100G |
| Fetal Bovine Serum | 5% | GeminiBio | 100-106 |
| PBS | 1X | prepared on-site | N/A |
| **Blocking solution 2** |  |  |  |
| Bovine Serum Albumin | 5% | Sigma | A2153-100G |
| Fetal Bovine Serum | 5% | GeminiBio | 100-106 |
| PBS | 1X | prepared on-site | N/A |

**Table 2:**
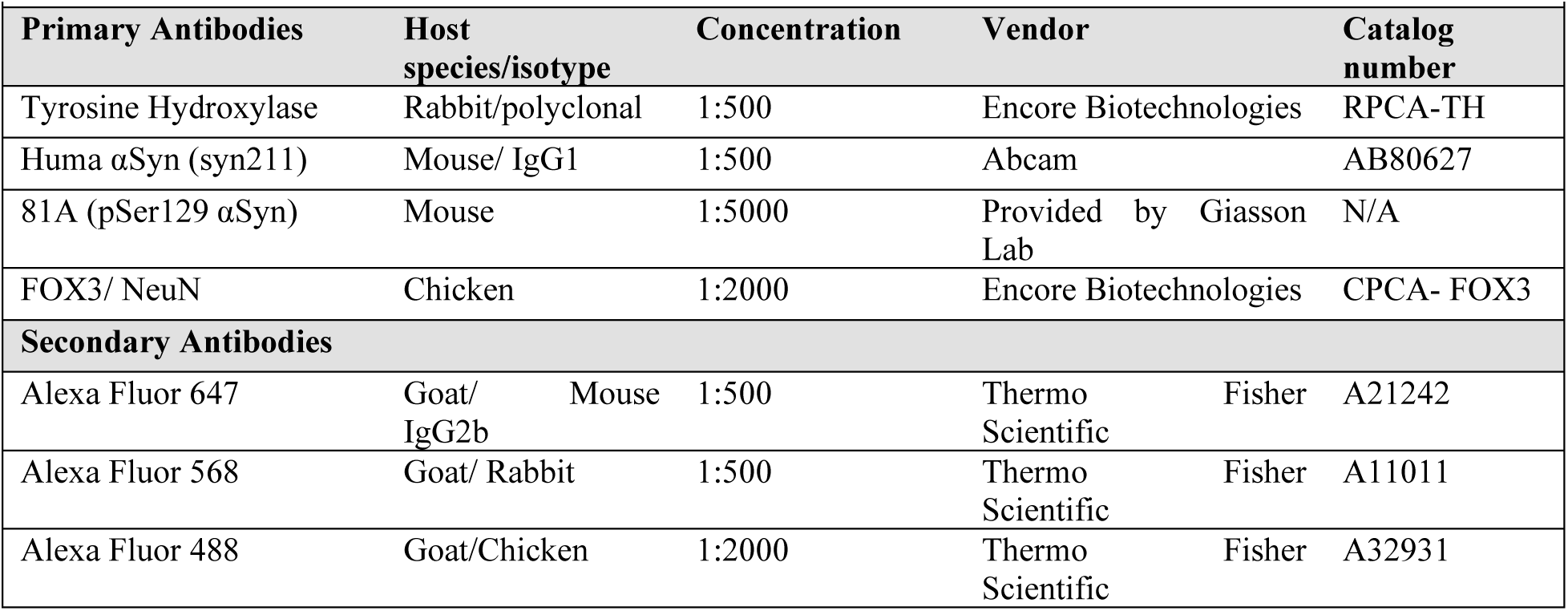
Antibody list.

### TUNEL Assay

The Elabscience® One-step TUNEL In Situ Apoptosis Kit (E-CK-A32 series) was used to detect apoptosis in tissue samples, following the manufacturer’s instructions. Tissue samples were mounted on gelatin-coated slides. A positive control was prepared by inducing DNA strand breaks with DNase I to verify the assay’s effectiveness and reagent performance. Briefly, 100 µL of 1× DNase I Buffer was added to the tissue sections and incubated at room temperature for 5 minutes. This was followed by the addition of 100 µL of DNase I solution (200 U/mL), which was incubated at 37°C for 10 - 30 minutes. After treatment, the sections were washed three times with PBS for 5 minutes each. The positive control was processed alongside experimental samples, using the same labeling and staining protocol to ensure procedural consistency. All tissue sections, including controls, were washed with PBS and permeabilized with 1× Proteinase K working solution for 20 minutes at 37°C. For labeling, slides were incubated with Terminal Deoxynucleotidyl Transferase (TdT) equilibration buffer, followed by the addition of a labeling solution containing TdT enzyme and fluorescein-labeled deoxyuridine triphosphate (dUTP). Samples were incubated at 37°C for 60 minutes in a humidified, light-protected chamber. Slides were then washed with PBS and counterstained with DAPI for nuclear visualization. Images were captured using a Nikon A1 laser-scanning confocal microscope (Nikon Instruments, Melville, NY) equipped with Plan Fluor 20× 0.75 NA and 40× 1.3 NA objectives. Z-stack images were acquired at 1 µm intervals across the entire specimen, and maximum intensity projection images were generated.

### Preparation of acute midbrain slices for single neuron recording and calcium imaging

Mice were perfused with ice-cold sucrose-sodium replacement solution containing (in mM): 250 sucrose, 26 NaHCO3, 2 KCl, 1.2 NaH2PO4, 11 dextrose, 7 MgCl2, and 0.5 CaCl2 (Table 1). Following perfusion, the mice were decapitated, and their brains were rapidly removed. For sectioning, the brains were cut rostrally to the prefrontal cortex (PFC) in the coronal plane to create a flat surface for attachment to the cutting stage. Coronal midbrain slices (250 µm thick), containing the substantia nigra and the ventral tegmental area, were prepared using a vibratome (VT1200S, Leica). The slices were transferred to a recovery chamber containing oxygenated artificial cerebrospinal fluid (ACSF) at 32°C. The ACSF composition was (in mM): 126 NaCl, 2.5 KCl, 2 CaCl2, 26 NaHCO3, 1.25 NaH2PO4, 2 MgSO4, and 10 dextrose, equilibrated with 95% O2 and 5% CO2. To minimize glutamate-induced toxicity during recovery, 10 µM MK-801 was added to the ACSF. The osmolarity of both the sucrose-sodium replacement solution and ACSF was adjusted to 300–310 mOsm, and the pH was maintained at 7.35–7.4. Slices were allowed to recover in the chamber for at least 30 minutes at 32°C before use in subsequent ex vivo electrophysiological and calcium imaging experiments.

### Electrophysiological Recordings and Analysis

Acute midbrain slices were transferred to a low-profile open diamond bath chamber (RC-26GLP, Warner Instruments, Hamden, CT, USA) and continuously perfused with oxygenated ACSF at 36–37°C. Dopamine neuron cell bodies were identified based on GCaMP6f fluorescence, using a 40× water immersion objective mounted on an Eclipse FN1 upright microscope (Nikon Instruments, Melville, NY, USA). The microscope was equipped with a Spectra X light engine (Lumencor, Beaverton, OR, USA) and a 470/24 nm solid-state illumination source. Images were captured using a 12-bit Zyla 4.2 sCMOS camera (Andor Technology, Belfast, Northern Ireland). Neuronal morphology was visualized with infrared differential interference contrast (IR-DIC) imaging. Borosilicate glass capillaries (1.5 mm O.D.; Sutter Instrument Company, Novato, CA, USA) were pulled into electrodes with a tip resistance of 4-8 MΩ using a P-2000 laser puller. The electrodes were filled with a potassium gluconate-based internal solution containing (in mM): 120 K-gluconate, 20 KCl, 2 MgCl2, 10 HEPES, 0.1 EGTA, 2 Na2ATP, and 0.25 NaGTP, with osmolarity adjusted to 290–295 mOsm and pH set to 7.25–7.30. Electrophysiological recordings were performed using an Axon Axopatch 200B microelectrode amplifier and digitized via a Digidata 1440A system using ClampEx 10.2 software (Molecular Devices, San Jose, CA, USA). Baseline recordings were obtained, followed by a hyperpolarization protocol as described in (Miller et al. 2021). During this protocol, baseline spontaneous firing activity was recorded for 30 seconds without current injection, followed by 30 seconds of hyperpolarization via injected current, and a subsequent 30-second recovery period to observe spontaneous firing activity. Data analysis was performed using Origin software, enabling precise quantification of neuronal responses under experimental conditions.

### Calcium Imaging and video processing

For calcium imaging experiments, AAV-syn-FLEX-jGCaMP8f-WPRE (1.1 x 10˄13U/µL), a gift from GENIE Project, 0.3 µL was injected into was injected into the SNc (ML = + 1.4 mm; AP = - 3.07 mm; DV = - 4.5 mm) and the VTA (ML = +/-0.4 mm; AP =-3.4 mm; DV =-4.4 mm) of DAT-Cre mice. Acute coronal slices containing the VTA and SNc were prepared and illuminated as previously described in (Miller et al. 2021). Images were acquired at a frame rate of 20–25 frames per second with no inter-frame delay. Baseline recordings were collected for 250 seconds for each experiment. Calcium imaging videos were processed by setting intensity thresholds to the observed minimum and maximum values for each video. Videos were converted to 8-bit TIFF format using ImageJ, and subsequently down-sampled for analysis. Trace extraction was performed using Miniscope 1-Photon-Based Calcium Imaging Signal Extraction Pipeline (MIN1PIPE), a semi-automated cell detection and activity quantification software, as referenced in (Lu et al. 2018)

### Network connectivity analysis and network graph construction

Network connectivity was assessed using pairwise Spearman’s rank correlation coefficients (MATLAB function: corr type: Spearman). A significance threshold of *p < 0.05* was applied, and networks with fewer than four neurons were excluded from the analysis. All network analyses utilized the weighted function implementation from the Brain Connectivity Toolbox. In all analyses, a “node” represents an individual neuron, and the normalized network degree is defined as the sum of all connections divided by the total number of neurons in the network, as previously described (Miller et al. 2021, Rubinov and Sporns 2010). Clustering coefficient, global efficiency, and network density are previously described in (Rubinov and Sporns 2010). Network graphs were constructed using undirected network connections. Data were imported into Gephi v0.9.2 for visualization.

### Statistical analysis

Statistical analyses were performed using MATLAB (version 2020a, MathWorks, Cambridge, MA, USA) and GraphPad Prism. Two-sample *t*-tests or one-way ANOVA were conducted where appropriate, with significance set at an alpha level of 0.05.

### Data and Code Availability

Region of interest (ROI) fluorescence data, as well as processed functional connectivity and network metrics, are available in MATLAB MAT format. Custom MATLAB code, designed to run as compiled code on a high-performance computing cluster, is available upon request. Additionally, MATLAB Live Script versions of the code, including both raw and processed data, can be provided to ensure reproducibility and ease of use.

## Results

### Establishing a pre-degenerative model of synucleinopathy

To establish a pre-degenerative model of synucleinopathy, we characterized dopaminergic neuron survival and αSyn expression following unilateral microinjection of either human α-synuclein (hαSyn; 1 µL of AAV-TH-hαSyn) or preformed αSyn fibrils (PFFs; 2 µg/µL) into the ventral tegmental area (VTA) or substantia nigra pars compacta (SNc) (Fig. 1). At four weeks post-injection, a timepoint chosen to precede significant neuronal loss, we assessed neuronal viability in midbrain slices encompassing the VTA and SNc using TUNEL staining and immunohistochemistry. The positive control for the TUNEL assay exhibited clear DNA fragmentation, confirming assay validity. However, no detectable TUNEL-positive cells were observed in either the VTA or SNc of either hemisphere following hαSyn or PFF injection, indicating the absence of significant apoptosis (Fig. 1). Immunohistochemical analysis was also performed for tyrosine hydroxylase (TH), pan-neuronal marker FOX3, and either phosphorylated αSyn (pSer129 αSyn; antibody 81A) to detect aggregated forms of αSyn following PFF injection, or hαSyn (antibody Syn211) following AAV injection. While the non-injected SNc and VTA of the contralateral hemisphere showed no 81A or Syn211 immunoreactivity (Fig. 2C, E, G, I), the SNc of the injected hemisphere exhibited staining for hαSyn or punctate 81A staining, confirming successful AAV-mediated expression of hαSyn or the presence of aggregated αSyn from PFF seeding, respectively (Fig. 2D, H). Notably, despite identical injection parameters, the VTA showed minimal hαSyn or 81A immunoreactivity compared to the SNc (Fig. 2F, J). Quantification of TH-positive (TH^+^) neurons revealed no significant difference between the injected hemisphere and the contralateral non-injected hemisphere in either the SNc or VTA (Fig. 2K, M), although a non-significant trend towards reduced TH^+^ cell number was observed in PFF-injected animals (Fig. 2M). Furthermore, the number of FOX3^+^ neurons remained unchanged (Fig. 2L and 2N), indicating preserved dopaminergic neuron density. Collectively, these data demonstrate that neither hαSyn expression nor the presence of PFF induces measurable dopaminergic neuron loss four weeks post-injection, confirming that this experimental paradigm represents an early, pre-degenerative stage of synucleinopathy. Given the absence of overt neuronal loss, we next investigated whether hαSyn or PFF-induced αSyn deposition alters the intrinsic electrophysiological properties of VTA and SNc dopamine neurons at this pre-degenerative stage.

**Figure 1.**
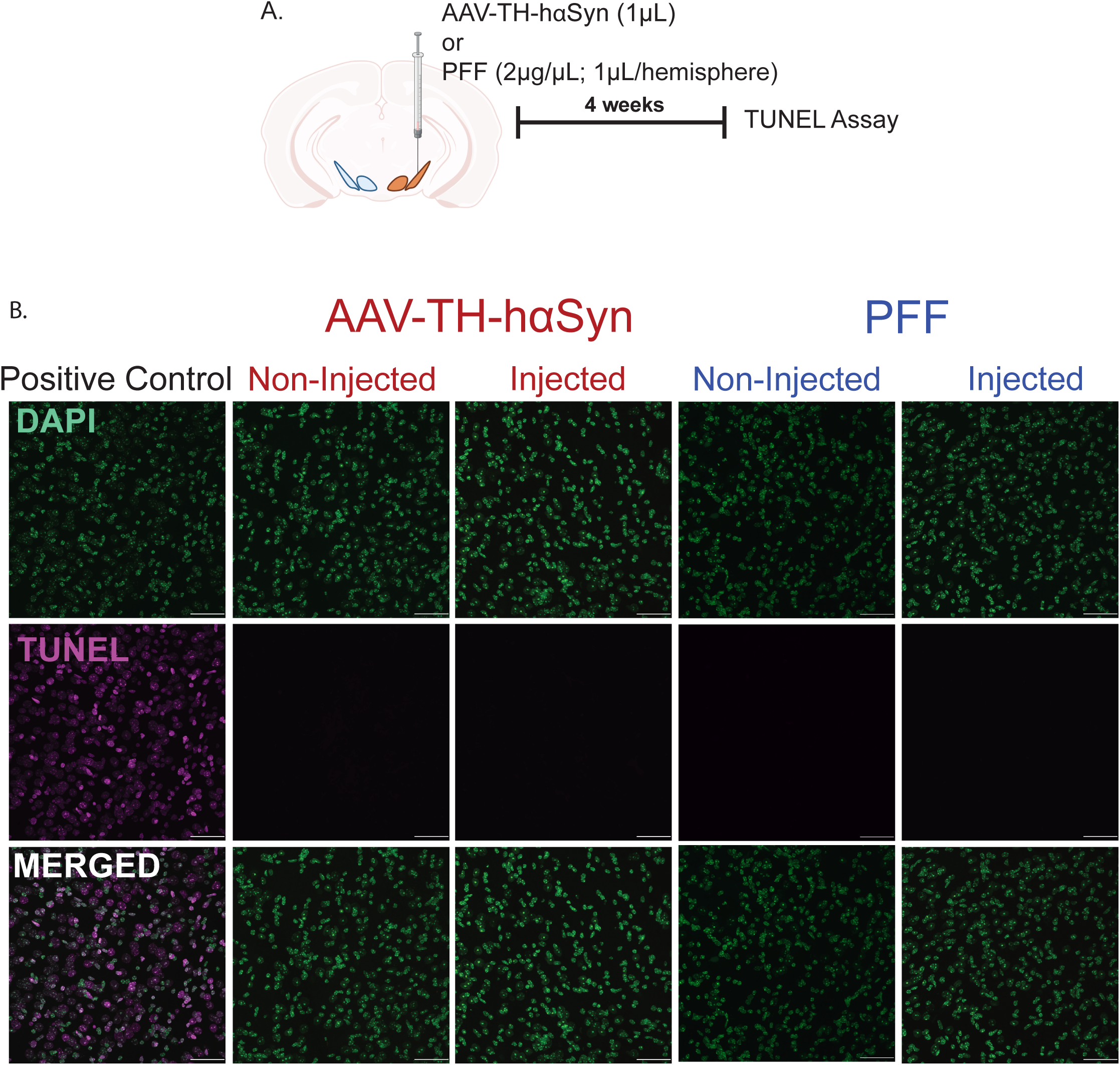
Assessment of neuronal viability in the VTA and SNc after AAV1-TH-hαSyn and PFF injections using TUNEL assay. **(A)** Schematic of the experimental design: Mice received injections of AAV1-TH-hαSyn (1 µL) or PFF (2 µg/µL) into the midbrain. Four weeks post-injection, brain sections were processed for TUNEL staining to assess apoptotic cell death. (**B**) Representative fluorescent images of DAPI^+^ nuclei and TUNEL^+^ cells in positive control and experimental sections. Midbrain sections from the contralateral (non-injected) and the ipsilateral (injected) sides of the AAV1-TH-hαSyn and PFF injection sites display DAPI^+^ nuclei but are entirely TUNEL-negative, indicating no detectable apoptotic cell death. This suggests that neurons in the VTA and SNc remain viable at this time point. Scale bar = 20 µm.

**Figure 2.**
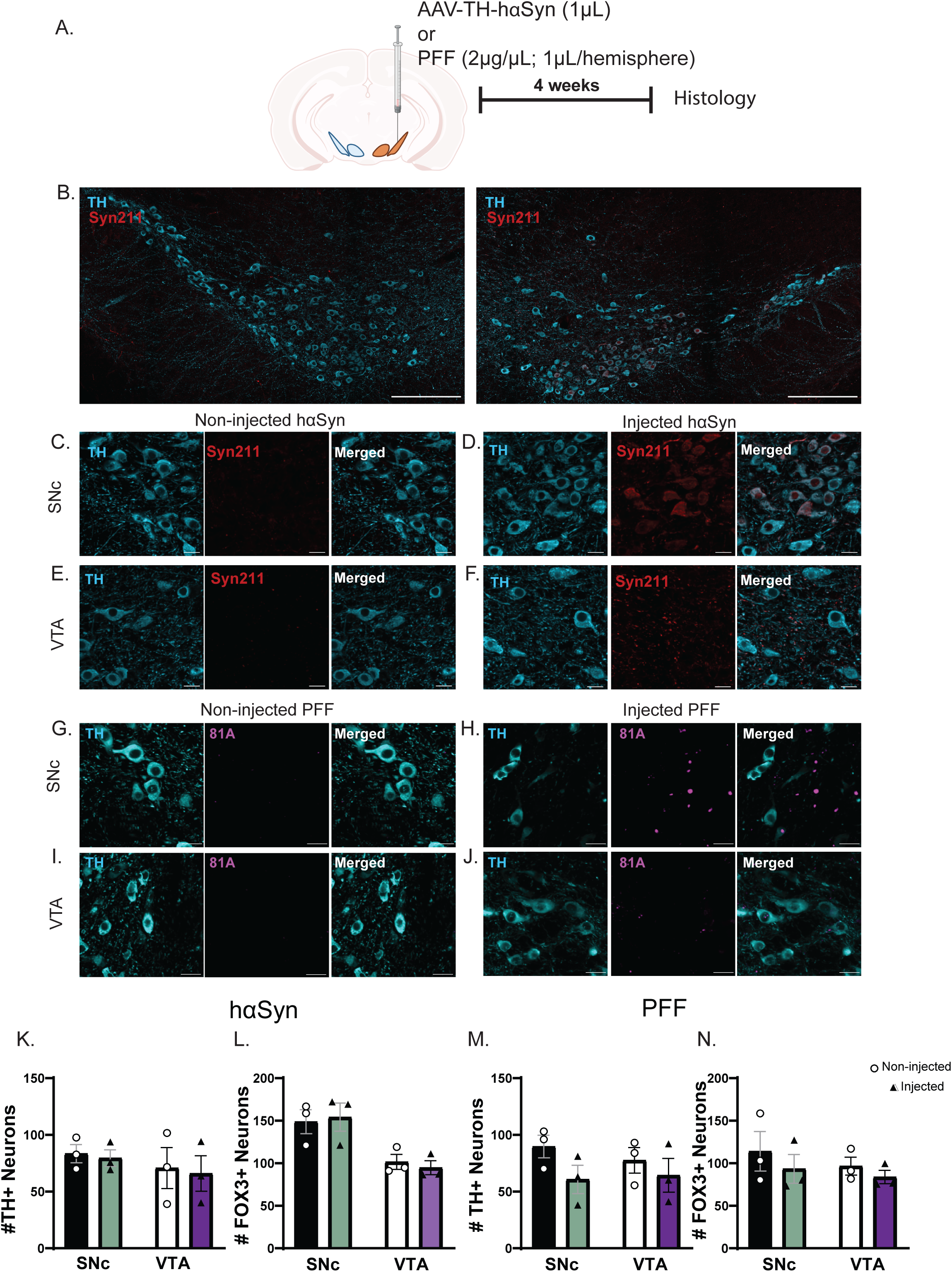
AAV-TH-hαSyn or pre-formed fibrils (PFF) increase corresponding immunostaining in the substantia nigra pars compacta (SNc) and ventral tegmental area (VTA). (**A)** Experimental design: Mice received unilateral injections of either AAV1-TH-hαSyn (1 µL) or PFF (2 µg/1 µL) into the midbrain. (**B)** Representative coronal section of the ventral midbrain (SNc and VTA), showing the injected (contralateral) and non-injected (ipsilateral) hemispheres. Four weeks post-injection, immunostaining with Syn211 (detecting hαSyn) shows colocalization in TH+ neurons on the injected side in the SNc **(D)** and VTA **(F)**, but not on the non-injected side respectively **(C, E)**. Following the same timeline, 81A (detecting phosphorylated αSyn aggregates) shows punctate signals localized within TH^+^ neurons on the injected side in the SNc **(H)** and VTA **(J)**, but not on the non-injected side respectively (**G, I)**. These data confirm the presence of hαSyn and aggregated αSyn in both the VTA and SNc. (n=3-4 biological replicates). (**K, M)** Four weeks after injection of either AAV1-TH-hαSyn or PFF, there was no significant change in the number of TH^+^ neurons in the SNc or VTA. (**L, N)** Similarly, no differences were number of FOX3^+^ neurons in the SNc or VTA. Error bars represent mean ± SEM. (n=3 independent biological replicates, one-way ANOVA and t-test, *p<0.05) Scale bars: 250 µm **(B);** 20 µm **(C-J)**.

### hαSyn increases basal firing activity of SNc dopamine neurons but not VTA dopamine neurons

Prior research indicates that SNc dopamine neurons are more vulnerable to neurodegeneration, while VTA dopamine neurons exhibit relative resilience (Damier et al. 1999, Lieberman et al. 2017, Halliday et al. 2005, Lebowitz and Khoshbouei 2020, Guzman et al. 2010). To investigate the impact of αSyn pathology (using hαSyn or PFF models) on these adjacent neuronal populations, we conducted whole-cell patch-clamp electrophysiology on midbrain slices (Fig. 3A, Fig. 4A). Spontaneous firing rates were measured in both the SNc and VTA regions from the injected ipsilateral and non-injected contralateral hemispheres (Fig. 3B-C, Fig. 4B-C). Single-neuron recordings examining hαSyn expression were performed four weeks following hαSyn expression (Fig. 3B-C). Consistent with previous studies (Zhang, Liu and Jiang 2021, Dagra et al. 2021, Lin et al. 2021), hαSyn expression in the SNc significantly increased the basal firing frequency of dopamine neurons on the injected side compared to the non-injected hemisphere (t-test, p < 0.05; Fig. 3D). Conversely, PFF-induced deposition did not alter the basal firing activity of SNc dopamine neurons (t-test, p > 0.05; Fig. 4D). Basal firing rates of VTA dopamine neurons remained unchanged by either haSyn or PFF (Figs. 3G, 4G). Furthermore, action potential morphology, including half-width and membrane capacitance, was unaltered in both SNc and VTA neurons (Figs. 3E-F, 3H-I, 4E-F, and 4H-I). This suggests that the hαSyn-induced increase in firing activity of SNc dopamine neurons does not stem from alterations in intrinsic membrane properties or cell size at four weeks post-hαSyn expression or PFF injection. These findings offer insights into the early pathophysiology of synucleinopathies, preceding neuronal degeneration. Specifically, hαSyn has been shown to increase calcium activity through D2 receptor and dopamine transporter (DAT)-mediated mechanisms (Dagra et al. 2021, Butler et al. 2015). However, the underlying mechanisms of PFF-induced dysregulation/degeneration remain unknown. Moreover, VTA neurons maintained stable firing patterns (Figs. 3G, 4G). While VTA dopamine neuron activity is preserved at 4 weeks post-hαSyn expression and PFF injection, prolonged hαSyn expression may eventually impair the function of these neurons. These results underscore the importance of exploring how synucleinopathies affect neuronal resilience, particularly in response to hyperpolarization-mediated stressors.

**Figure 3.**
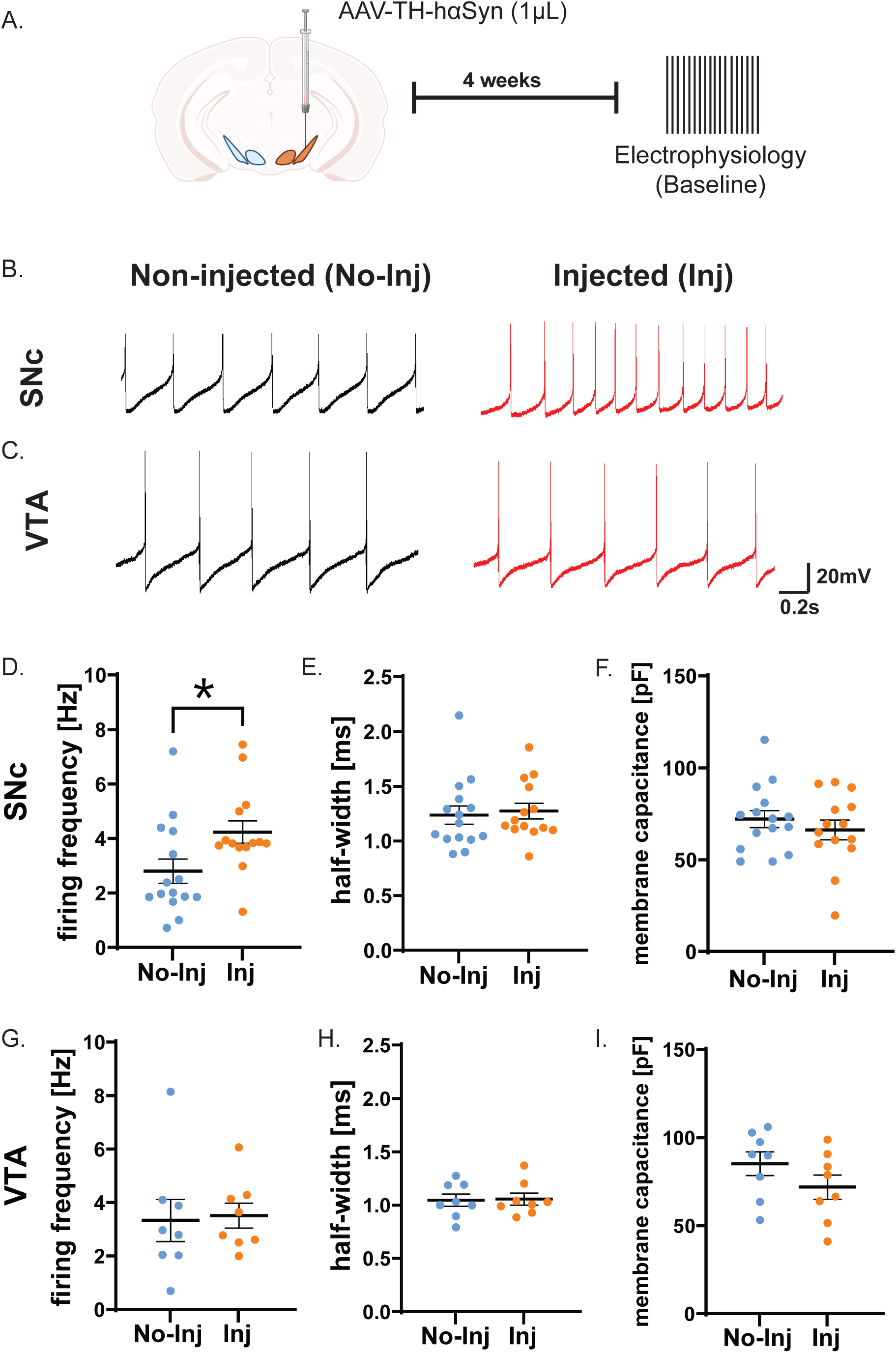
hαSyn alters the electrophysiological properties of dopamine neurons in the SNc but not in the VTA dopamine neurons. (**A**) Schematic representation of the experimental design for single-neuron recordings conducted four weeks post hαSyn injection (1 µL) into the midbrain. (**B**) Representative electrophysiological traces from SNc dopamine neurons in the non-injected and hαSyn-injected sides. (**D**) The basal firing frequency of SNc dopamine neurons is significantly increased on the ipsilateral side post-hαSyn injection. (**C, G**) Representative traces from VTA dopamine neurons on the contralateral and ipsilateral sides demonstrate that hαSyn injection does not alter the basal firing frequency of VTA dopamine neurons. (**E, F**) Bar graphs depicting the half-width and membrane capacitance of SNc dopamine neurons. (**H, I**) Bar graphs depicting the half-width and membrane capacitance of VTA dopamine neurons. Error bars represent the mean ± SEM. Statistical significance was determined using a t-test (*p < 0.05; n = 3-4 independent biological replicates).

**Figure 4.**
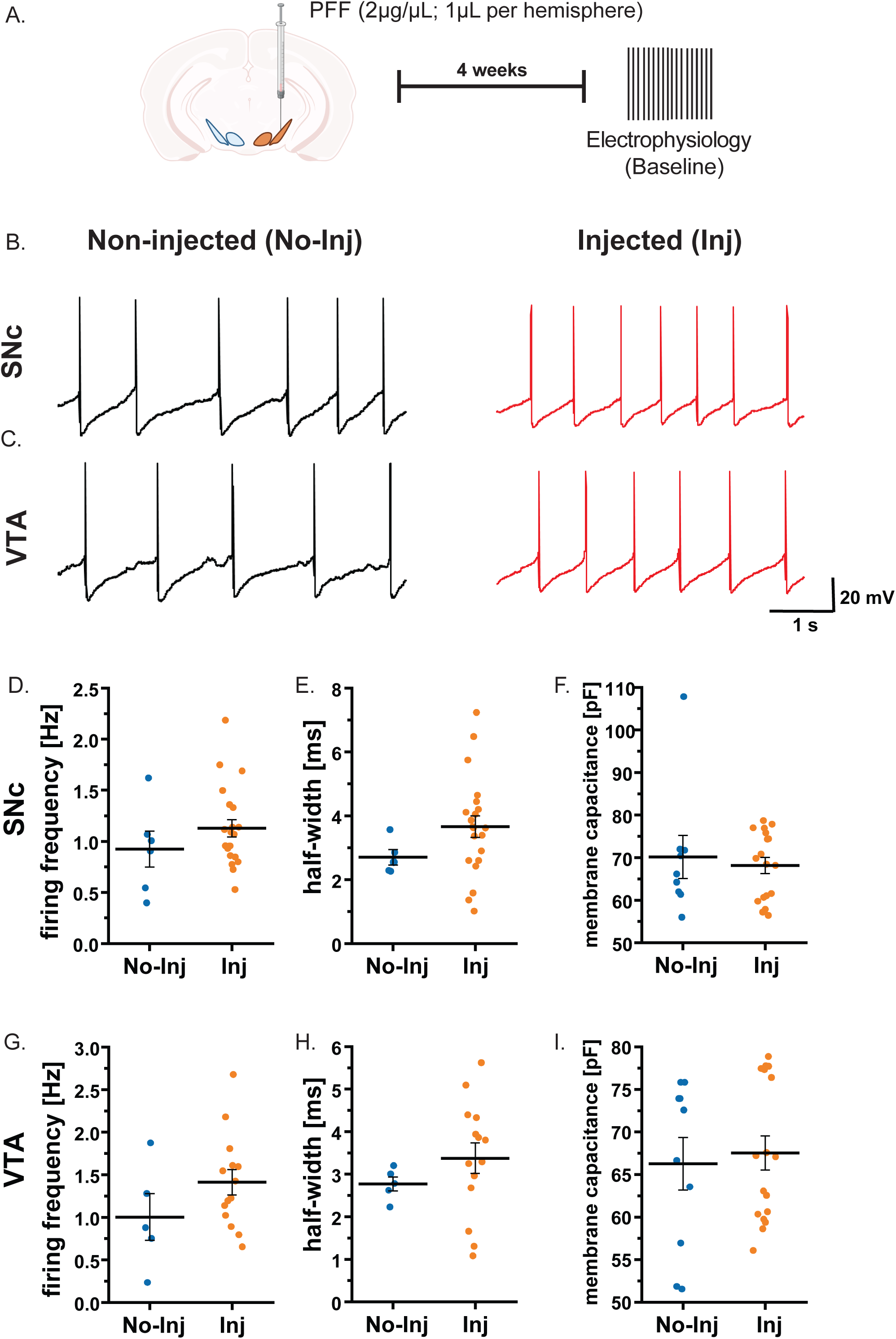
The presence of PFF does not change the electrophysiological properties of dopamine neurons in the SNc or VTA dopamine neurons. Schematic representation of the experimental design for single-neuron recordings conducted four weeks post-unilateral PFF deposition (2µg/µL) into the midbrain. (**B)** Representative electrophysiological traces from SNc dopamine neurons in the non-injected and PFF-injected sides. (**D**)The basal firing frequency of SNc dopamine neurons is similar in PFF-injected and non-injected mice. (**C, G)** Representative traces from VTA dopamine neurons in PFF-injected and non-injected mice demonstrate that PFF injection does not alter the basal firing frequency of VTA dopamine neurons. (**E, F, H, I**) Bar graphs depicting the half-width and membrane capacitance of SNc and VTA dopamine neurons. Error bars represent the mean ± SEM (4-6 independent biological replicates). Statistical significance was determined using a t-test.

### PFF and hαSyn increase SNc dopamine neuron vulnerability to hyperpolarization challenge, while VTA neurons remain resilient

Midbrain (SNc and VTA) dopamine neurons maintain spontaneous pacemaking activity, relying on finely tuned homeostatic mechanisms. To assess how hαSyn or PFF affects neuronal resilience under stress, we applied a hyperpolarization protocol (Miller et al., 2021) using whole-cell patch-clamp electrophysiology in slices from mice injected four weeks prior with AAV1-TH-hαSyn (Fig 5A), or PFF (Fig. 6A) in the midbrain. This approach allowed us to compare the pre-and post-hyperpolarization firing activity of SNc and VTA dopamine neurons from injected ipsilateral hemisphere and the non-injected contralateral hemisphere. Interestingly, SNc dopamine neurons on the side injected with either AAV1-TH-hαSyn (Fig 5), or PFF (Fig. 6) displayed significantly reduced firing rates following hyperpolarization-induced stress, while those on the contralateral non-injected hemisphere quickly recovered (Fig 5F and 6F). In contrast, VTA dopamine neurons demonstrated comparable resilience on both injected ipsilateral hemisphere and the non-injected contralateral hemisphere, maintaining stable firing rates after the hyperpolarization challenge (Fig. 5G and 6G). These results underscore the SNc’s susceptibility to αSyn pathology. Over time, this increased vulnerability may contribute to the loss of SNc dopamine neurons seen in PD. Together, our findings reveal that during pre-degenerative state, hαSyn expression selectively increase firing rates among SNc dopamine neurons without causing measurable cell death (Fig.3). SNc neurons are less capable of coping with acute perturbations in neuronal homeostasis, suggesting that these early functional disruptions could set the stage for neurodegeneration. In contrast, VTA neurons remain relatively unaffected at four weeks post-hαSyn expression or αSyn PFF deposition, supporting their reported resilience (Mosharov et al. 2009, Damier et al. 1999, Halliday et al. 2005). Understanding these initial functional changes provides a critical foundation for developing therapeutic strategies aimed at preserving dopaminergic function and preventing the progression of αSyn-associated pathology. In addition, although understanding hαSyn pathology at the single-neuron level is important, it is equally important to investigate how hαSyn pathology affects dopamine neuronal networks through their integrated activity. Therefore, to bridge the gap from single-neuron physiology to ensemble-neuronal network level, we next examined how hαSyn expression or PFF deposition influence network connectivity in the SNc and VTA.

**Figure 5.**
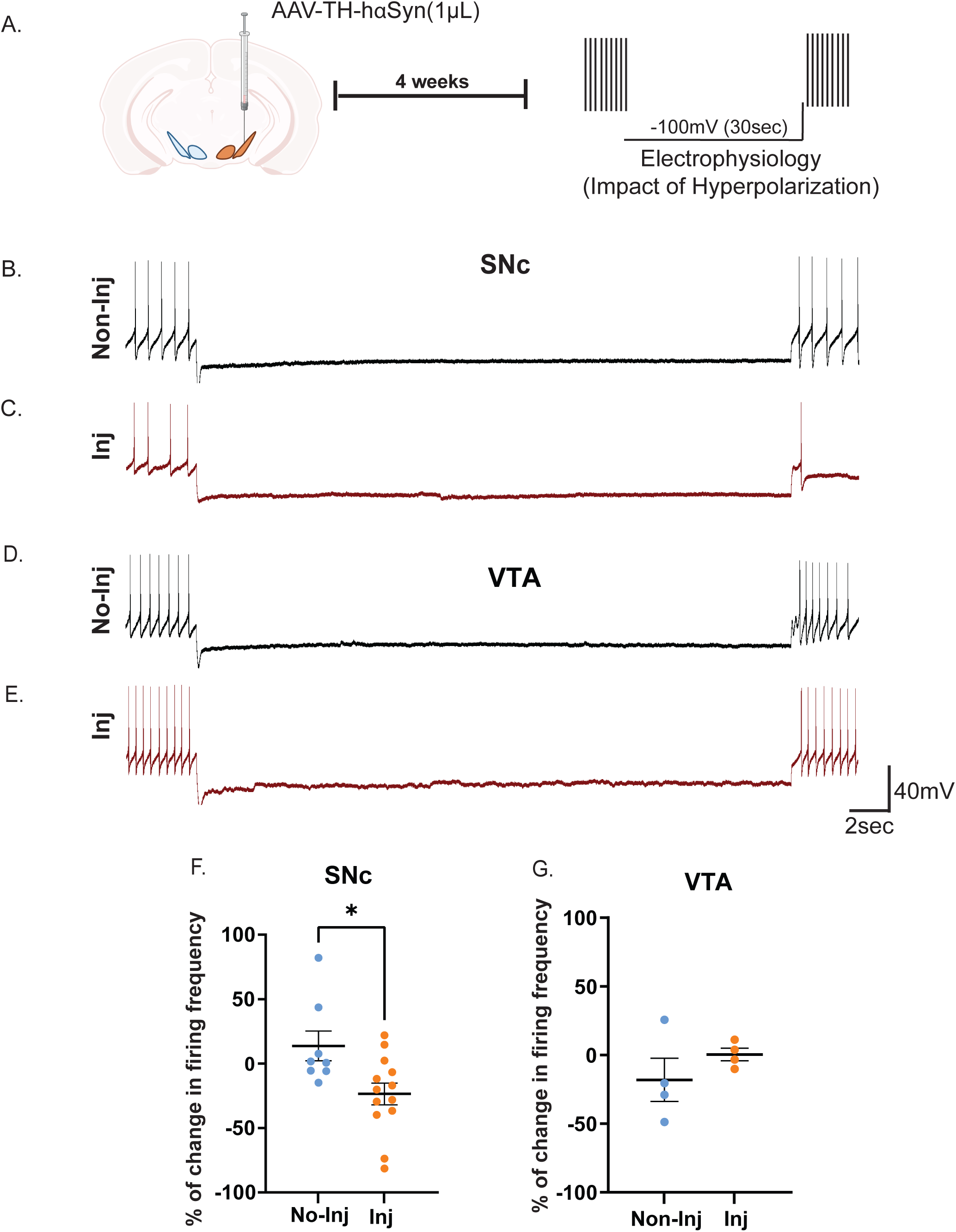
hαSyn overexpression compromises the resilience of SNc dopamine neurons to hyperpolarization challenge (acute stress). (**A**) Experimental design: Four weeks after unilateral hαSyn injection (1 µL), basal firing frequency of SNc dopamine neurons was recorded both ipsilaterally (injected side) and contralaterally (non-injected side). Neurons were then hyperpolarized to –100 mV for 30 seconds. Following restoration of the resting membrane potential, post stress firing frequency was assessed to determine resilience to hyperpolarization. **(B, C)** Representative electrophysiological traces from SNc dopamine neurons before and after the 30-second hyperpolarization step. (**F**) The accompanying graph shows the percentage change in firing frequency, revealing a diminished capacity of SNc neurons to recover following hyperpolarization stress in the presence of hαSyn. **(D, E)** Representative traces from VTA dopamine neurons at baseline and post-hyperpolarization. **(G)** The associated graph demonstrates that VTA neurons effectively buffer the hyperpolarization-induced perturbation, maintaining firing rates without significant changes after hαSyn overexpression. Error bars represent mean ± SEM (n = 3-4 independent biological replicates). Statistical significance was assessed by t-test (*p < 0.05).

**Figure 6.**
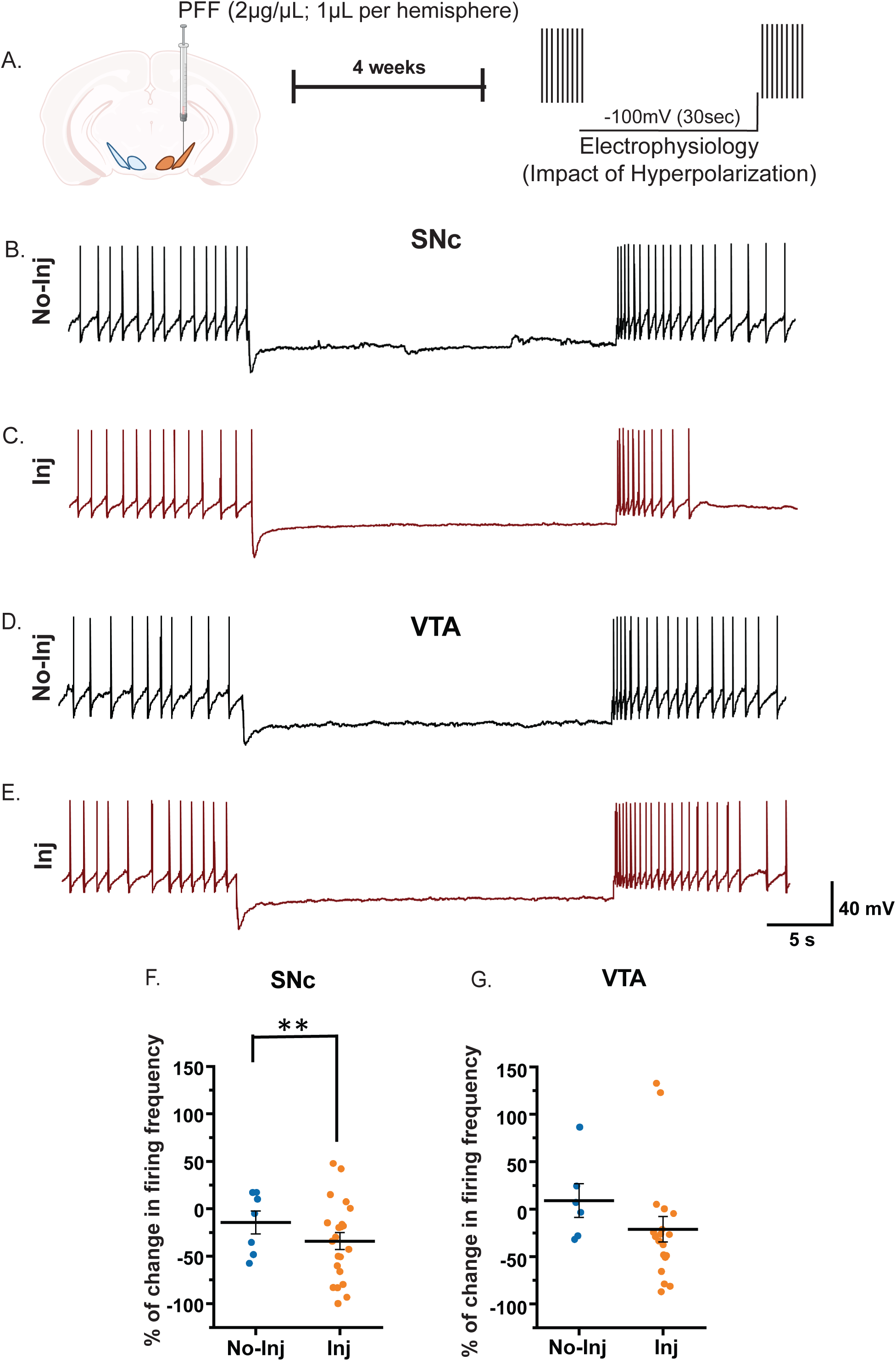
PFF deposition impairs the resilience of SNc dopamine neurons to hyperpolarization challenge (acute stress). **(A)** Experimental design: Four weeks post unilateral PFF deposition (2 µg/µL), the basal firing frequency of SNc dopamine neurons was recorded both ipsilaterally (injected side) and contralaterally (non-injected side). Neurons were then hyperpolarized to –100 mV for 30 seconds. After returning to resting membrane potential, post-stress firing frequency was measured to assess resilience. **(B, C)** Representative electrophysiological traces from SNc dopamine neurons before and after the 30-second hyperpolarization step. (**F**) The corresponding graph quantifies the percentage change in firing frequency, showing a reduced capacity of SNc neurons to recover following hyperpolarization stress when exposed to PFF. **(D, E)** Representative traces from VTA dopamine neurons at baseline and post-hyperpolarization. **(G)** The associated graph indicates that VTA neurons maintain stable firing frequencies and remain unaffected by PFF deposition under these conditions. Error bars represent mean ± SEM (n = 4–6 independent biological replicates). Statistical significance was determined by t-test (**p < 0.01).

### During the pre-degeneration phase, αSyn and PFF deposition increased network hyperconnectivity in the SNc, but not the VTA

To investigate the impact of PFF or hαSyn accumulation on dopamine network dynamics in the SNc and VTA, we injected DAT:cre mice with a combination of AAV5-FLEx-GCaMP8f (0.3µL. see Table 1) and either AAV1-TH-hαSyn or PFF into the SNc or VTA. GCaMP8f is a genetically encoded calcium indicator, serving as a surrogate for studying neuronal activity (Zhang et al. 2023). Changes in GCaMP8f fluorescence intensity provide a real-time, non-invasive approach to monitor the activity of individual neurons and neuronal populations. The fast response time of GCaMP8f allows for capturing rapid neuronal firing, improving the temporal resolution. Post-PFF or AAV1-TH-hαSyn-injection, midbrain slices were prepared and imaged to monitor GCaMP8f fluorescence signal as an indicator of neuronal activity (Fig. 7, A–C). Calcium imaging videos were processed using MIN1PIPE (Miniscope 1-Photon–Based Calcium Imaging Signal Extraction Pipeline; see Methods), which extracts neural signals and identifies regions of interest (ROIs) with spatial footprints. From these data, calcium traces were isolated for individual neurons (Fig. 7C), and raster plots of neuronal activity were generated for the SNc and VTA, binned at 100-second intervals (Fig. 7D). All traces were then correlated using Spearman’s rank correlation, applying a significance threshold of *p* < 0.05 for network analysis. Neurons (nodes) with significant connections are labeled “1” in yellow, whereas nodes with self-connections or non-significant connections are labeled “0” in green (Fig. 7E), with the corresponding network graph shown in Fig. 7F.

**Figure 7.**
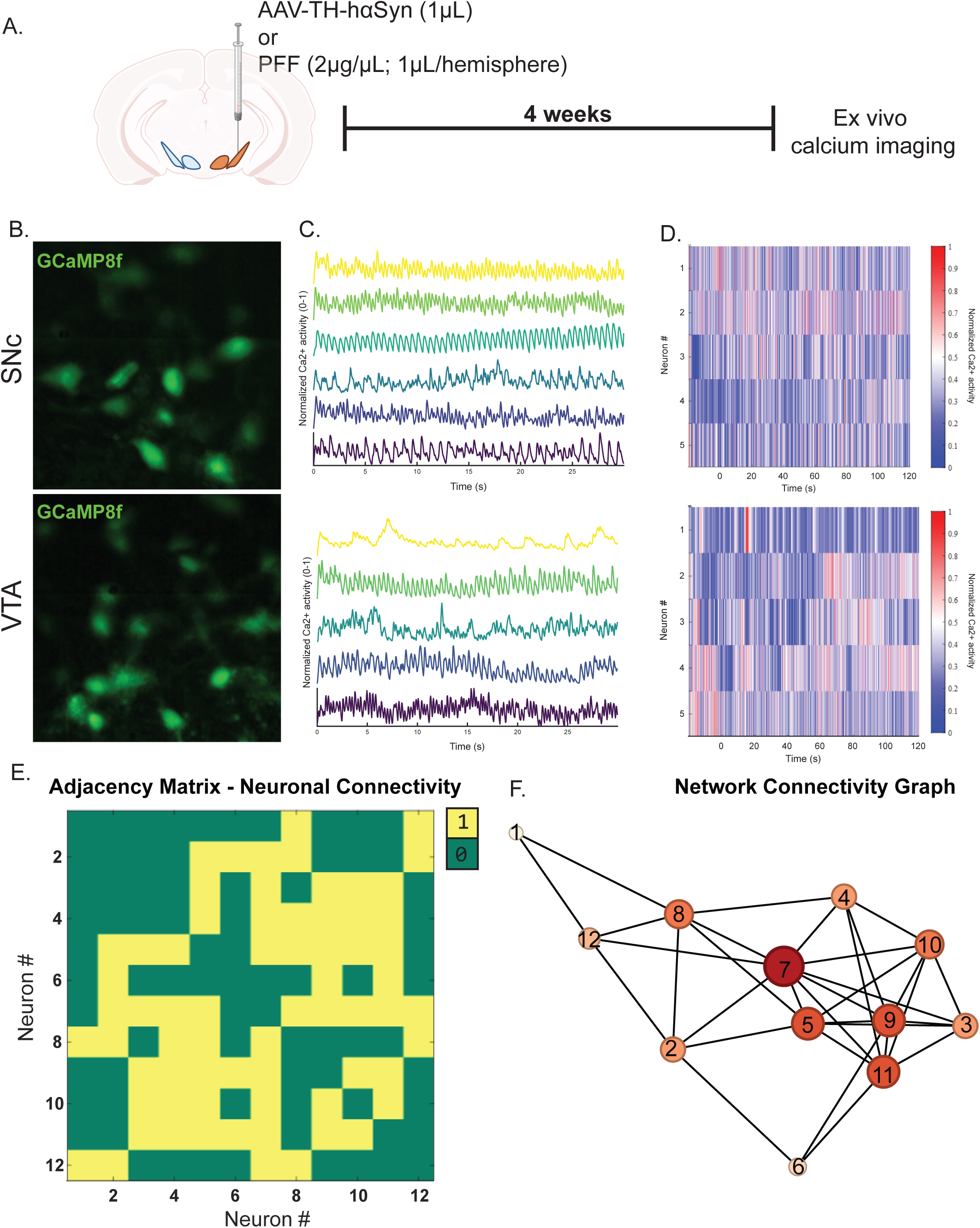
Investigating hαSyn-and PFF-induced regulation of dopamine neuronal network connectivity in the SNc and VTA. (**A**) Schematic representation of the experimental design: DAT:cre mice were bilaterally injected with AVV5:FLEx-GCaMP8f (300nL) for live cell calcium imaging and unilaterally received AAV-TH-hαSyn (1 µL) or PFF (2µg/µL). (**B**) Representative GCaMP8f-labeled dopamine neurons. (**C**) Normalized calcium traces from dopamine neurons in the SNc (top) and VTA (bottom). (**D**) Raster plots of neuronal firing activity show temporal patterns of calcium signaling in individual dopamine neurons. (**E**) Adjacency matrices illustrating neuronal connectivity, where “1” denotes a connection between two neurons and “0” denotes no connection. (**F**) Network connectivity graphs derived from adjacency matrices. Each node represents an individual neuron, with the size and intensity of color indicating the number of connections for that neuron (greater size, larger number of connections and darker color reflect higher connectivity). These analyses highlight the differential network connectivity of dopamine neurons in the SNc and VTA under conditions of hαSyn or PFF exposure.

To confirm that the measured neuronal activity stemmed from fluctuations in the fluorescent signals of individually isolated neurons within the network, we examined the relationship between inter-neuron distance and correlated activity in all cells. Across all experimental conditions (hαSyn and PFF, both injected hemisphere and non-injected hemisphere) in the SNc and VTA, we observed no relationship between distance and correlation (Supplemental Fig. 1, A–G). This finding suggests that changes in network connectivity parameters were neither due to fluorescent bleed-over from nearby neurons nor neuronal death (inactivity). Among the network connectivity parameters measured were node degree, clustering coefficient, network density, and global efficiency (Yin et al. 2020, Rubinov and Sporns 2010).

Node degree describes the number of links (interconnections) a node has with other nodes. Since the total number of connections is influenced by the total number of neurons in the region of interest, node degree is normalized by dividing by the total number of nodes, yielding a normalized node degree. This metric represents the average number of connections per neuron. Consistent with our electrophysiological data, both hαSyn and PFF enhanced network connectivity (node degree) among SNc dopamine neurons relative to control conditions. Specifically, SNc dopamine neurons injected with AAV1-TH-hαSyn or PFF exhibited higher normalized node degree distributions, as well as increased average normalized node degree compared to controls (*p*<0.001 and *p*<0.05, Fig. 8, A–C, E–F). In contrast, VTA network connectivity measures (normalized node degree) remained unchanged (Fig. 8, D, G–H). These findings suggest that hαSyn or PFF accumulation leads to a hyperconnected dopamine neuronal network specifically in the SNc, correlating with the increased firing rates observed both in the present study and in previous reports (Lin et al. 2021, Dagra et al. 2021). Thus, prior to cell death in synucleinopathies, SNc dopamine neurons bear a heightened physiological burden, evident at both the single-neuron and network levels.

**Figure 8.**
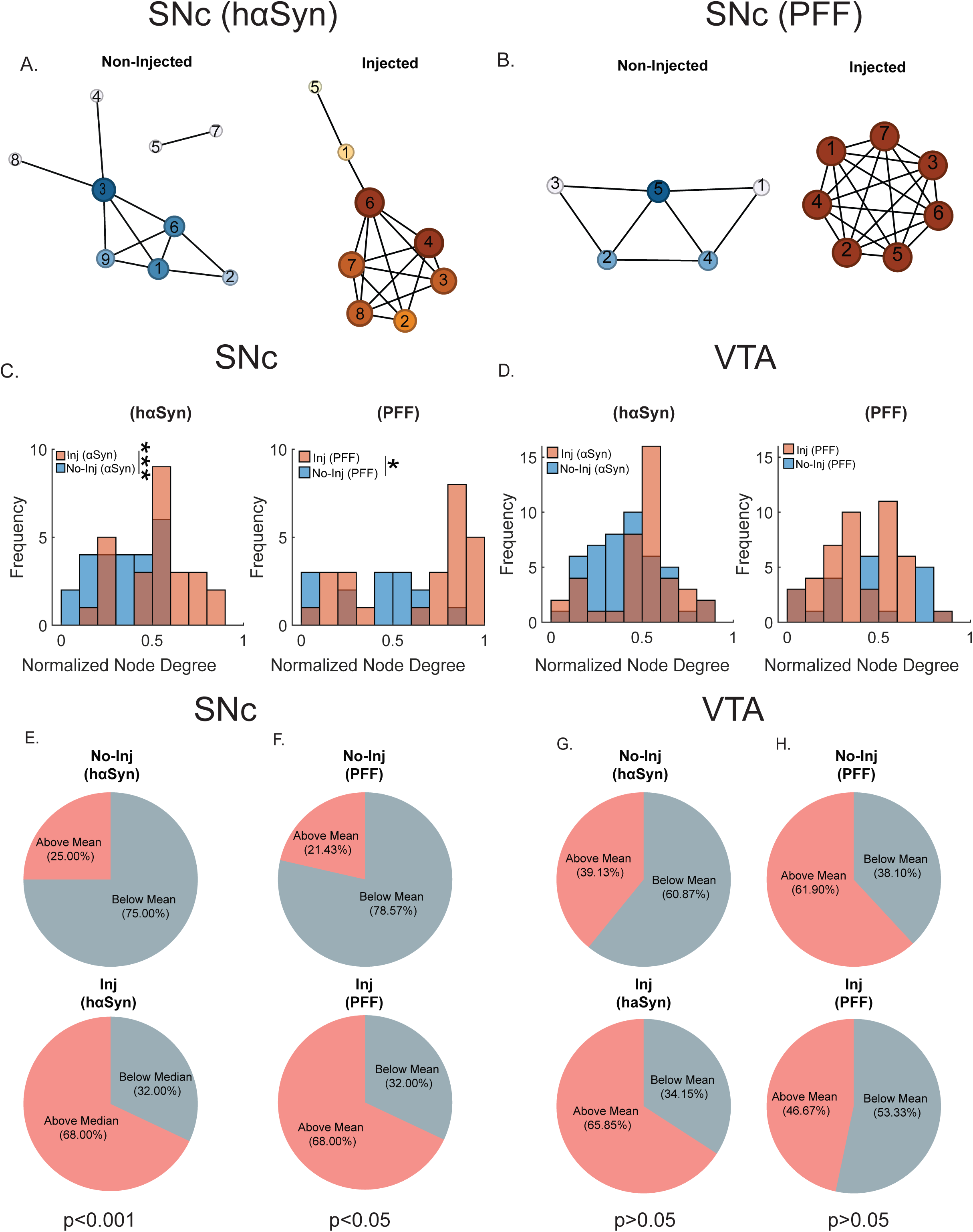
Both hαSyn expression and PFF deposition increase hyperconnectivity in the SNc, but not the VTA. **(A)** Left panel: Network graphs of the SNc for non-injected and the hαSyn-injected hemispheres (AAV-TH-hαSyn, 1 µL). (**B**) Network graphs of the SNc for non-injected and the PFF-injected hemispheres (PFF, 2µg/µL). **(C)** Bar graphs of the frequencies of normalized node strengths in non-injected versus hαSyn-or PFF-injected SNc. (**D**) Bar graphs of the frequencies of normalized node degree in non-injected versus hαSyn-or PFF-injected VTA. (**E-H**) Pie graphs of the frequencies of the node strength above the mean of non-injected and hαSyn-or PFF-injected side in the SNc and VTA. (n = 3-5 independent biological replicates, t-test, * p<0.05, ***p<0.001).

We also examined higher-order parameters of network metrics, namely clustering coefficient, network density, and global efficiency (Supplemental Fig. 2). The clustering coefficient is a measure of functional segregation that quantifies the proportion of triangular connections around each node. A high clustering coefficient indicates networks in which neurons share many mutual neighbors (i.e., subpopulations of highly interconnected nodes), whereas a lower value indicates a more decentralized or random structure. We observed no change in the mean clustering coefficient following hαSyn or PFF injection in either the SNc or VTA (Supplemental Fig. 2A) indicating that there was a similar level of highly interconnected subpopulations.

Network density measures the fraction of existing connections out of all possible connections within a network, and global efficiency refers to the average inverse shortest path length— an index of how readily information travels across the network. Higher global efficiency indicates the potential for parallel information processing and better communication between distant nodes. Notably, only hαSyn deposition increased network density and global efficiency in the SNc (but not in the VTA; *p*<0.05, t-test, Supplemental Fig. 2, B–C). These parameters remained unchanged after PFF deposition in both the SNc and VTA (Supplemental Fig. 2, B–C).

Consistent with single-neuron recordings, hαSyn significantly increased node degree, network density, global efficiency, and spontaneous firing in dopamine neurons. Conversely, PFF only increased node degree in the SNc; other network properties and basal firing remained unchanged. These findings highlight that hαSyn expression has a substantially greater impact four weeks post-injection on basal firing and network properties compared to PFF deposition. It is important to note that we encountered significant challenges when patch-clamping dopamine neurons with PFF deposition, suggesting a potential imminent failure state. Thus, prolonged PFF deposition might lead to a significant reduction in both network density and global efficiency. Collectively, these findings indicate that αSyn accumulation drives early alterations in neuronal and network activity prior to neurodegeneration.

## Discussion

This study provides new insights into the differential early-stage vulnerability of dopaminergic neurons in the SNc compared to those in the VTA during αSyn-associated pathology. Although mounting evidence implicates αSyn aggregates in the selective degeneration of SNc neurons characteristic of PD, the mechanisms triggering their early neuronal dysfunction remain poorly understood. In the current study, by employing PFF or viral-mediated hαSyn overexpression (AAV-TH-hαSyn) before overt cell death, we delineated how αSyn pathobiology disrupts the physiological activity and network connectivity of dopaminergic neurons in the SNc and VTA.

Several key findings emerge from our work. First, we observed that four weeks postinjection, alteration in αSyn (hαSyn or PFF) did not induce measurable loss of dopamine neurons in either the SNc or VTA. This allowed us to capture a “pre-degenerative” stage, one in which functional deficits precede structural collapse. At this early stage, we found that although hαSyn expression and PFF deposition occurred in the midbrain, SNc dopamine neurons increased spontaneous firing rates and neuronal network hyperconnectivity. Such changes were absent in the VTA dopamine neurons, highlighting a regional difference. Critically, we also show that SNc dopamine neurons become less resilient to acute stress (hyperpolarization challenge), suggesting that αSyn-mediated alterations render these neurons more vulnerable to subsequent insults.

These electrophysiological and network-level perturbations in SNc dopamine neurons before cell death resonates with prior studies. SNc neurons are known for their large, complex axonal arbors and an intrinsic pacemaking phenotype reliant on L-type calcium channels, metabolic homeostasis, and sustained bioenergetic demands (Pacelli et al. 2015, Braak et al. 2003b, Guzman et al. 2010, Geibl et al. 2024, Sulzer 2007, Surmeier, Obeso and Halliday 2017). By contrast, VTA dopamine neurons, though similar in many respects, express distinct molecular markers (e.g., calbindin) and operate under different metabolic and synaptic constraints (Mosharov et al. 2009, Fu et al. 2012, Dopeso-Reyes et al. 2014, Lammel, Lim and Malenka 2014, Lieberman et al. 2017). These interregional differences have been proposed as contributing factors to the selective vulnerability observed in the SNc dopamine neurons in PD. Secondly, αSyn can bind to several membrane proteins such as Na^+^, K^+^-ATPase, Na^+^ channels, and K^+^ channels. αSyn has been shown to increase basal Na+ levels, preventing return to baseline membrane potential in neurons (Shrivastava et al. 2015).

Our data align with the hypothesis that αSyn aggregates, even before outright neuronal degeneration, can affect neuronal activity in a manner that promotes instability at both the single-neuron and network levels. The known roles of αSyn in synaptic vesicle trafficking and neurotransmitter release could be altered by its pathological forms, leading to dysregulated dopamine handling, increased basal extracellular dopamine, and an overall hyper-excitable state (Butler et al. 2015, Kramer and Schulz-Schaeffer 2007, Stoker and Greenland 2018). Indeed, previous work suggests that αSyn triplication or overexpression leads to increased baseline excitability and altered dopamine dynamics (Swant et al. 2011, Butler et al. 2015, Miller et al. 2021) potentially increasing the SNc neurons’ energetic and synaptic stress. The increased neuronal activity and network hyperconnectivity may tax cellular homeostatic mechanisms, including mitochondrial function and calcium buffering, thereby setting the stage for neuronal demise.

In contrast, VTA dopamine neurons in our models remained stable, showing no significant changes in firing properties or network connectivity, at least at four weeks following PFF or hαSyn accumulation. This resilience may reflect intrinsic physiological differences between these dopamine neuron populations. VTA dopamine neurons are less reliant on the same calcium handling pathways as SNc dopamine neurons and may retain more robust adaptive responses to stress or αSyn burden. Furthermore, the distinct molecular signatures of VTA dopamine neurons might confer protective advantages, such as differential αSyn clearance mechanisms, or synaptic plasticity profiles that limit the impact of pathological αSyn forms.

Our network connectivity analyses reveal that hαSyn or PFF accumulation induce a more synchronized, hyperconnected environment in SNc, but not VTA, dopamine neurons. Enhanced neuronal connectivity could arise from augmented synaptic coupling or altered inhibitory-excitatory balances, ultimately rendering the SNc microcircuit more vulnerable to disruptions. The shift to a hyperconnected state may make the network less capable of compensating for perturbations, like metabolic stress. As αSyn aggregates spread and escalate in complexity, this fragile network may fail under sustained burden, leading to progressive synaptic dysfunction and loss of dopamine neurons. These results have important implications for understanding the early pathogenic events in PD. Identifying a pre-degenerative window in which neuronal dysfunction is present and could be reversible (Dagra et al. 2021, Lin et al. 2021) offers a crucial time frame for therapeutic intervention. Targeting αSyn aggregation, modulating firing rates, or normalizing network connectivity during these early stages may prevent or delay the cascade of events leading to irreversible loss of dopamine neurons. Additionally, exploring why VTA dopamine neurons resist these early changes may help identify protective mechanisms that could be harnessed to bolster SNc dopamine neurons against αSyn pathology.

In conclusion, our study underscores that before cell death, αSyn pathobiology differentially affects the SNc and VTA dopamine neurons. By disrupting firing properties and network stability selectively in SNc dopamine neurons, αSyn sets the stage for later neurodegeneration. Future work should explore the cellular and molecular underpinnings of this differential vulnerability, including metabolic demands, and synaptic mechanisms that could be leveraged to halt or slow disease progression.

---

**Supplemental Figure 1.**
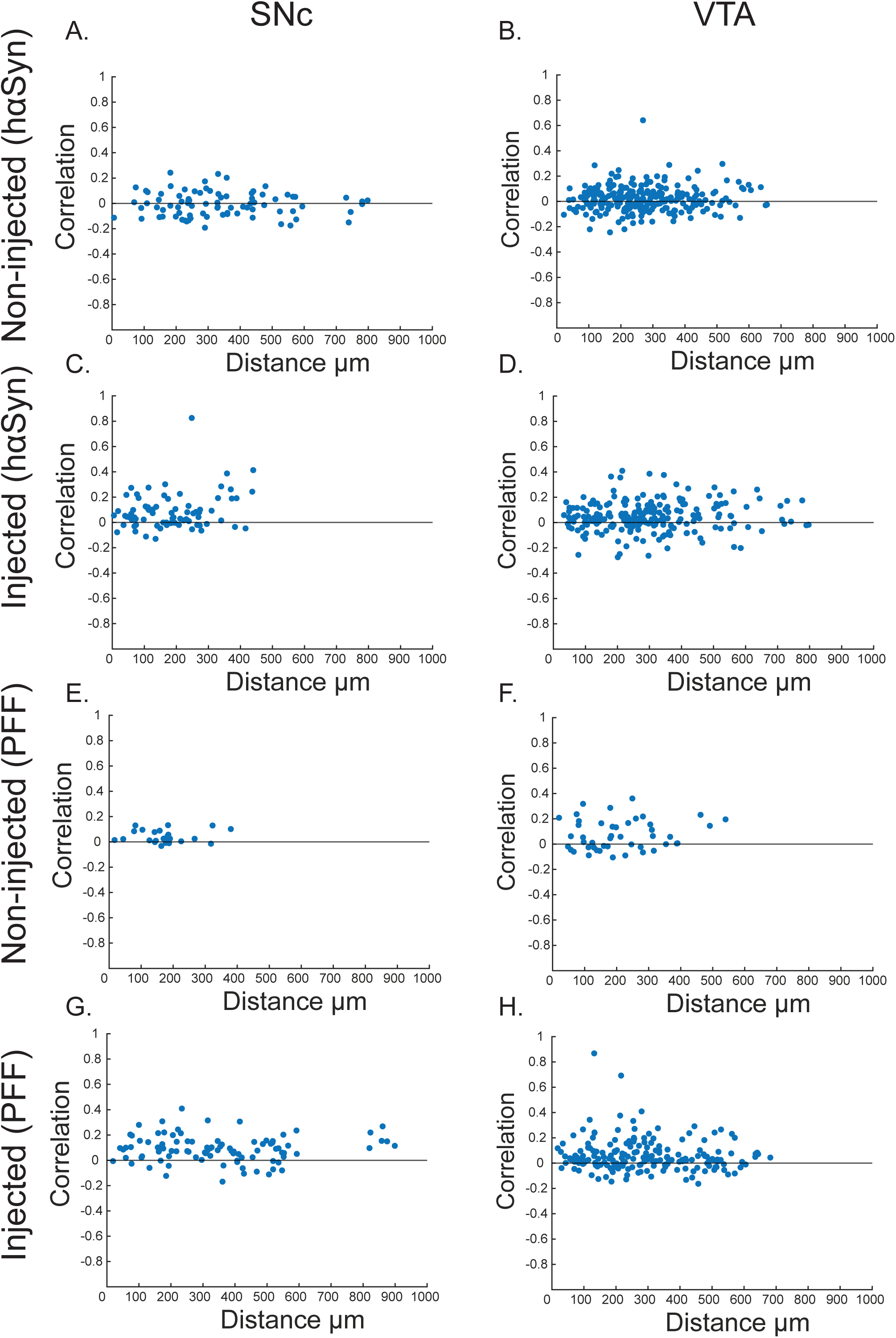
Neuronal correlation activity does not correlate with inter-cell distance. Plots illustrate the relationship between correlation values and inter-neuron distance in the SNc for the non-injected hαSyn hemisphere (**A**), injected hαSyn (**C**), non-injected PFF hemisphere (**E**), and injected PFF hemisphere (**G**). Corresponding plots for the VTA are shown for the non-injected hαSyn hemisphere (**B**), injected hαSyn (**D**), non-injected PFF hemisphere (**F**), and injected PFF hemisphere (**H**).

**Supplemental Figure 2.**
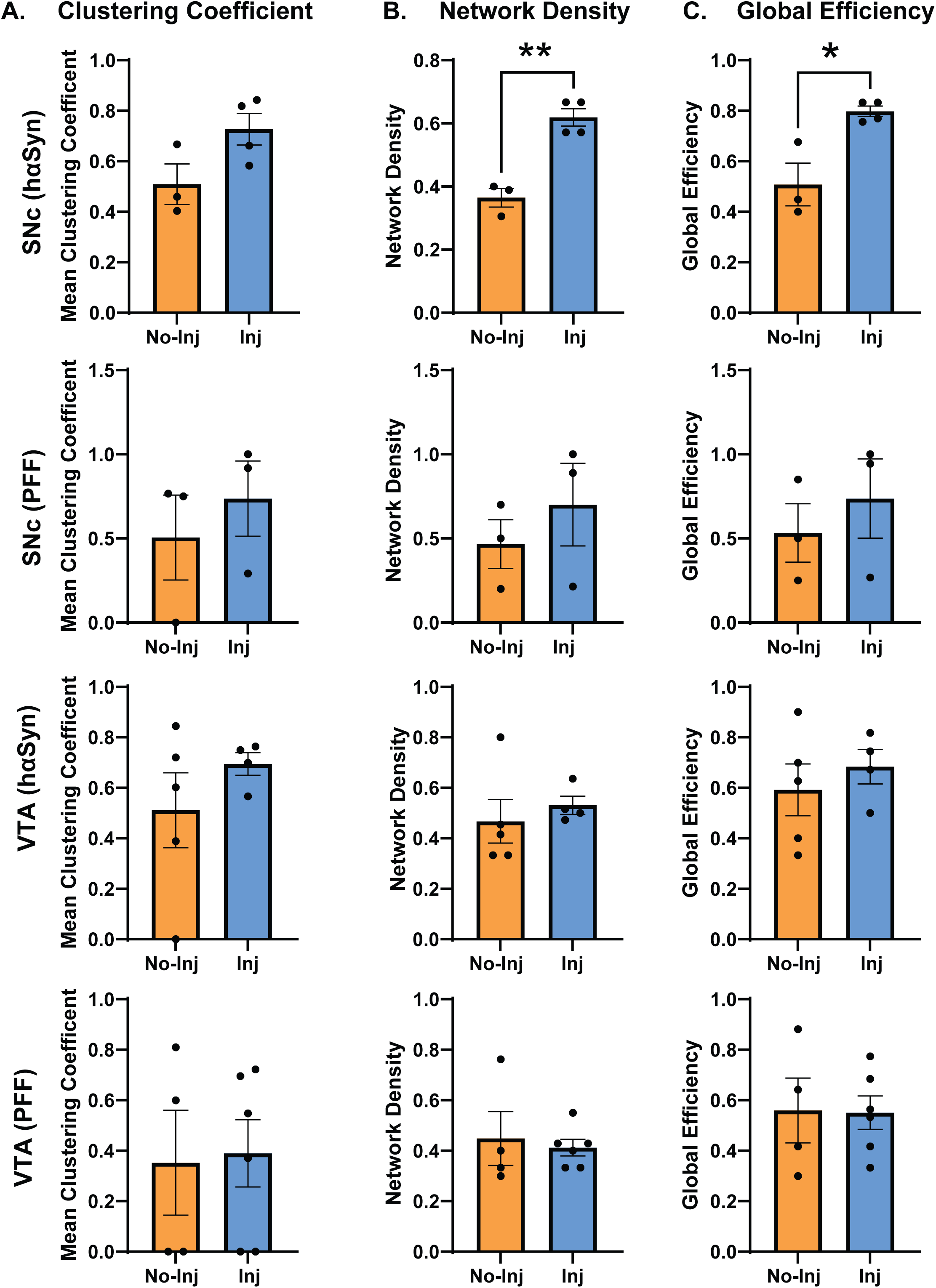
hαSyn increases global network indices but not subnetwork parameters. (**A**) Mean clustering coefficient in hαSyn-or PFF-injected, compared to the non-injected hemisphere. (**B**) Network density in hαSyn-or PFF-injected, compared to the non-injected hemisphere in the SNc and VTA. (**C**) Global efficiency in hαSyn-or PFF-injected, compared to the non-injected hemisphere in the SNc and VTA (n = 3-5 independent biological replicates, t-test, * p<0.05, **p<0.01).

